# When does heterogeneity average out in coarse-grained ecological communities?

**DOI:** 10.64898/2026.10.09.757735

**Authors:** Mingyu Luo, Van Savage, Chuliang Song

## Abstract

Aggregating components into a few variables, such as species into total biomass, is common in complex systems, yet differences among components can still shape aggregate dynamics. For dynamics near a stable equilibrium, we quantify this irreducible dynamical heterogeneity as the spectral mismatch between the aggregate’s fluctuations and those of the best closed aggregate model, an Ornstein–Uhlenbeck process of the same dimension. In weakly interacting random communities, increasing species richness can reduce or increase this heterogeneity, depending on interaction structure. Our framework clarifies when ecological and evolutionary differences average out and when their dynamical effects persist.

## Introduction

Ecology is often forced to speak in aggregates. Long time series may record total biomass but not species composition; demographic models may lump age or stage structure into a few classes; spatial surveys may report regional abundance while local patch states remain unresolved [Fig. 1(a)]. Such aggregates have long been central in ecosystem modeling [1–3] and are routinely treated as dynamical states, fitted with ecological models, and used to infer resilience, stability, and responses to perturbation [4–6]. Treating such an aggregate as an exact dynamical state requires closure: two communities with the same aggregate must have the same expected aggregate change, regardless of hidden species composition, patch composition, or stage structure.

**FIG. 1.**
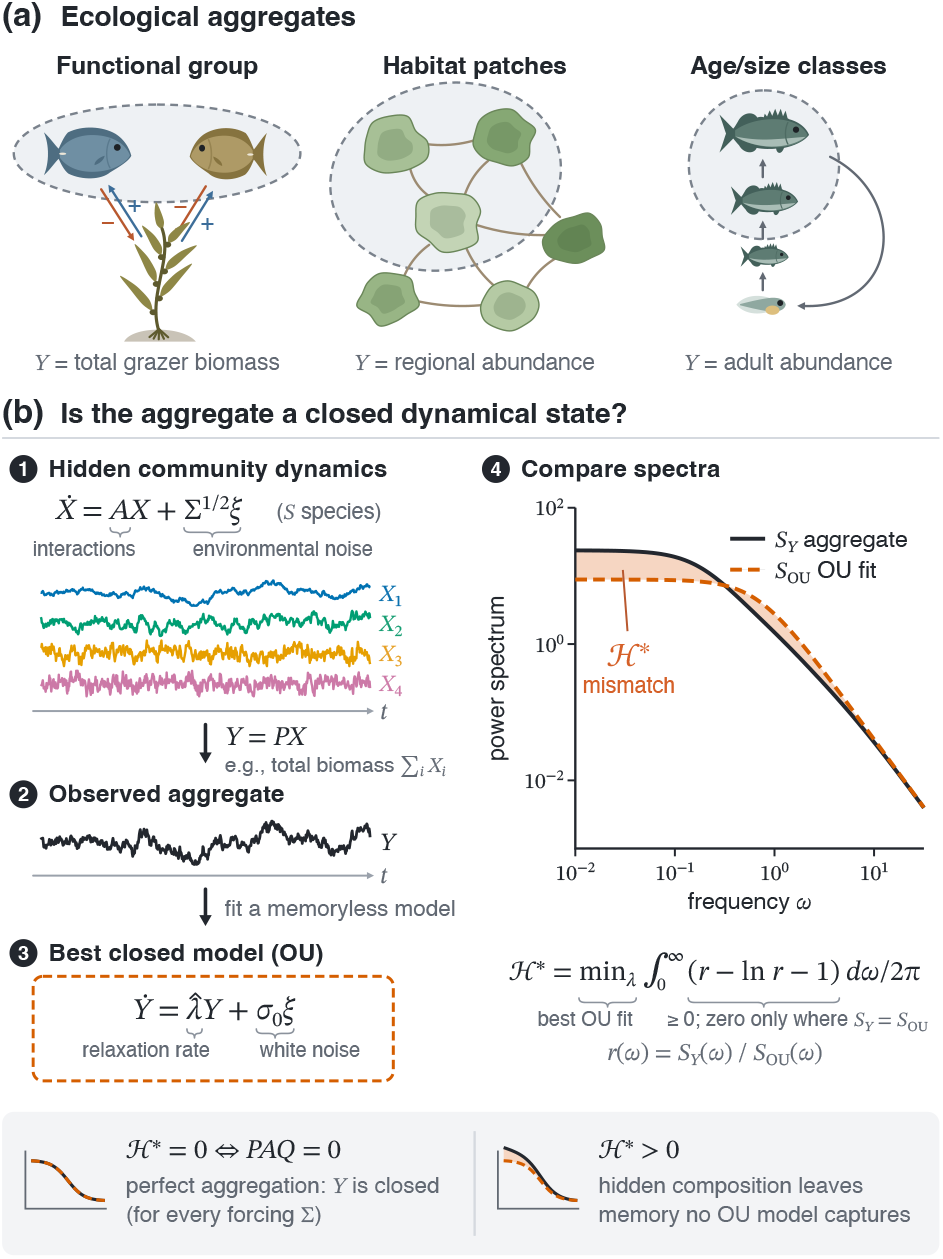
Coarse-graining and spectral closure. (a) Three ecological aggregates: a functional group (two congeneric grazers feeding on kelp; arrows denote positive and negative effects), a region of habitat patches linked by dispersal (shading indicates carrying capacity), and adult stage classes of a size-structured population. Dashed boundaries enclose the grouped variables; *Y* is the resulting aggregate. (b) The community state *X*, governed by 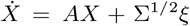, is observed only through *Y* = *PX* (steps 1–2). The best closed Ornstein– Uhlenbeck model is fitted to *Y* (step 3), and its spectrum *S*_OU_(*ω*) is compared with the exact spectrum *S*_*Y*_ (*ω*) (step 4). The Itakura–Saito divergence between the two, minimized over the OU rate, defines the irreducible dynamical heterogeneity ℋ* (shaded). ℋ* vanishes for every Σ *≻* 0 if and only if *PAQ* = 0. Example: four weakly competing species with self-regulation rates 0.2, 1, 3, and 6, Σ = *I*, and total biomass as the aggregate (ℋ \**≈* 0.09); time series are standardized for display.

The classical formalization of this closure question is the perfect aggregation framework of Iwasa, Andreasen, and Levin [7], building on related aggregation ideas in economics and control theory [8, 9]. Let *X*∈ ℝ^*n*^ be the full, microscopic state and *Y* = *PX*∈ℝ^*m*^ the aggregate, or macroscopic, state, where *m < n* and the projection matrix *P* ∈ℝ^*m×n*^ has full row rank and defines the aggregation. Perfect aggregation asks whether *Y* has autonomous dynamics exactly consistent with the full dynamics. For linearized dynamics 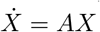, with community Jacobian *A*, the condition becomes *PAP* ^+^*P* = *PA*, where *P* ^+^ is the Moore–Penrose pseudoinverse. Equivalently, with *Q* = *I−P* ^+^*P* the orthogonal projection onto ker(*P*), the hidden subspace of changes that leave *Y* unchanged,

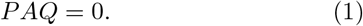

Thus hidden variation at fixed aggregate has no effect on the aggregate derivative. For total biomass, *P* = **1**^*⊤*^, where **1** is the all-ones vector, this requires all column sums of *A* to be equal: every species must have the same instantaneous net effect on total biomass.

Perfect aggregation is a clean but knife-edge condition. Approximate aggregation theory instead chooses an inconsistency criterion and finds the aggregate model that minimizes it [10–12]. Different criteria preserve different objects—instantaneous vector fields, finite-time predictions, temporal averages, or trajectories—so the optimal aggregate model depends on the modeling goal. Here we compare stationary fluctuation spectra and quantify the unavoidable error when an aggregate is represented by a closed, same-dimensional Ornstein–Uhlenbeck (OU) model. Related information-theoretic work quantifies whether microscopic dynamics provide predictive information beyond a macroscopic process [13]; here the aggregate is prescribed and the benchmark is specifically an OU process of the same dimension.

Here we develop a theoretical framework for quantifying the dynamical effects of ecological heterogeneity after aggregation. Its metric, ℋ*, is the spectral mismatch remaining after fitting the best same-dimensional OU aggregate model.

### Irreducible dynamical heterogeneity

We now formalize the aggregation question as a stationary spectral comparison [Fig. 1(b)]. Let *X* denote small deviations from a stable equilibrium, and let

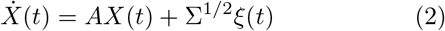

be the linearized stochastic community dynamics. Throughout, *A* is Hurwitz stable, Σ *≻* 0 is the covariance of environmental forcing, and *ξ*(*t*) is Gaussian white noise with *⟨ξ*_*i*_(*t*)*ξ*_*j*_(*t*′)*⟩* = *δ*_*ij*_*δ*(*t − t*′).

For a scalar aggregate *Y* = *p*^*⊤*^*X* with nonzero weight vector *p ∈* Rℝ^*n*^, the stable system has a unique stationary Gaussian solution. The stationary spectrum *S*_*Y*_ (*ω*) describes how the variance of the aggregate is distributed across angular frequencies *ω*, where *ω >* 0 corresponds to a fluctuation period 2*π/ω*. With the usual two-sided spectral density, restricted here to *ω ≥* 0, it is

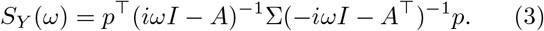

For total biomass, *p* = **1**. Equation (3) gives the aggregate response to hidden modes through the full-system resolvent and avoids assumptions about projected unresolved dynamics.

The simplest closed, memoryless model for a scalar aggregate is a one-dimensional Ornstein–Uhlenbeck process, the continuous-time analogue of a first-order autoregressive [AR(1)] model,

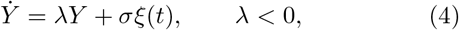

where *−* 1*/λ* is the relaxation time and *σ*^2^ is the noise intensity. Its spectrum is

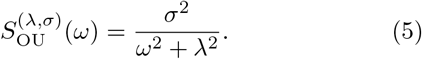

We measure the departure from this benchmark with the Itakura–Saito spectral divergence [14],

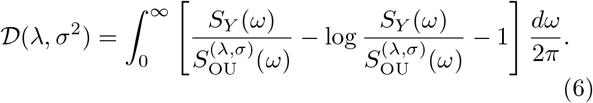

The integrand has the form *x −* log *x−*1 *≥*0, with equality only at *x* = 1, so *D* = 0 if and only if the exact and OU spectra coincide at all frequencies. For stationary Gaussian fluctuations, *D* is the expected loss of Whittle log-likelihood per unit time incurred by modeling the aggregate with the OU spectrum [15]; equivalently, it is the Kullback–Leibler divergence rate between the aggregate process and the OU model [16, 17].

At high frequencies, the deterministic drift *AX* contributes only lower-order corrections, so *S*_*Y*_ (*ω*) ~ *p*^*⊤*^Σ*p/ω*^2^. Finite divergence requires the OU spectrum to share this tail, fixing the noise intensity at 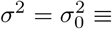 *p*^*⊤*^Σ*p* and leaving only the rate to be minimized. The unique minimizing rate is

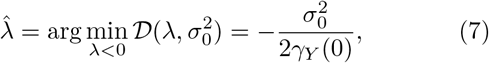

where *γ*_*Y*_ (*τ*) = *⟨Y* (*t*+*τ*)*Y* (*t*)*⟩* is the stationary autocovariance and *γ*_*Y*_ (0) is the aggregate variance. The fit matches the aggregate variance and initial autocorrelation decay rate, 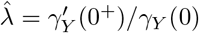 (Appendix A).

We define the irreducible dynamical heterogeneity of the scalar aggregate as this minimized spectral likelihood loss,

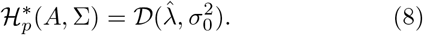

Because the divergence is integrated over frequency, 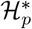 has units of inverse time. For total biomass, we write 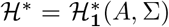. Unless otherwise stated, the examples use isotropic forcing, Σ = *I*. Duplicating a community into two isolated copies with independent forcing leaves ℋ* for their combined total biomass unchanged: both the aggregate spectrum and its optimal OU spectrum double. The same construction applies to vector aggregates *Y* = *PX∈* ℝ^*m*^, such as trophic compartments or spatial regions, with *P* of full row rank. The scalar spectrum is replaced by the *m×m* cross-spectral matrix of *Y*, and the benchmark by an *m*-dimensional OU process. This comparison includes both fluctuations of individual aggregates and temporal relationships between them. The high-frequency tail fixes the diffusion matrix of the OU model at *D*_0_ = *P* Σ*P* ^*⊤*^. Let *D*_*P*_ (*L*) denote the matrix extension of the scalar spectral loss for an OU model with stable drift *L*; its definition is given in Appendix A.

Let Γ =*⟨X*(*t*)*X*(*t*)^*⊤*^ *⟩* be the stationary community covariance, satisfying *A*Γ + Γ*A*^*⊤*^ + Σ = 0. The unique optimal drift is 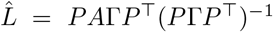. We define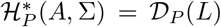; see Appendix A and the Supplemental Material [18] for the derivation.

For one specified positive-definite 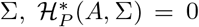 means that the stationary marginal Gaussian process *Y* has the same law as an *m*-dimensional OU process. It need not imply that the instantaneous aggregate drift *PAX* is determined by *PX*, because the microscopic state may retain additional predictive information. A structural condition independent of forcing is recovered by requiring vanishing for every positive-definite forcing covariance:

### Proposition

Assume *A* is Hurwitz stable. For a coarsegraining *Y* = *PX*, with *P*∈ ℝ^*m×n*^ of rank *m*,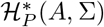 vanishes for every positive-definite Σ if and only if

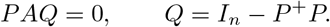

The condition must hold for every Σ: for a particular forcing covariance, a hidden mode can leave no trace in the aggregate power spectrum, giving 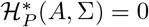 even though *PAQ* = 0 (the Supplemental Material gives an example with Σ = *I*); other forcing covariances generally reveal it. The proof is given in Appendix B. For a scalar aggregate *Y* = *p*^*⊤*^*X*, the condition reduces to *p*^*⊤*^ being a left eigenvector of *A*; for total biomass (*p* = **1**), this is the equality of all column sums of *A*, the perfect aggregation condition in Eq. (1).

Because ℋ* is a functional of the aggregate spectrum, it can in principle be estimated from sufficiently long, sufficiently resolved aggregate observations. A fully empirical estimate additionally requires estimating the aggregate spectrum and treating sampling and observation error; a discrete-time AR(1) formulation and a sampled-data benchmark are given in the Supplemental Material [18].

### Ecological examples

Three coarse-grainings illustrate how hidden variables become dynamically visible in an aggregate [Fig. 2]. In each case, the spectral mismatch remaining after optimal OU fitting quantifies how strongly ecological differences remain expressed in aggregate fluctuations.

**FIG. 2.**
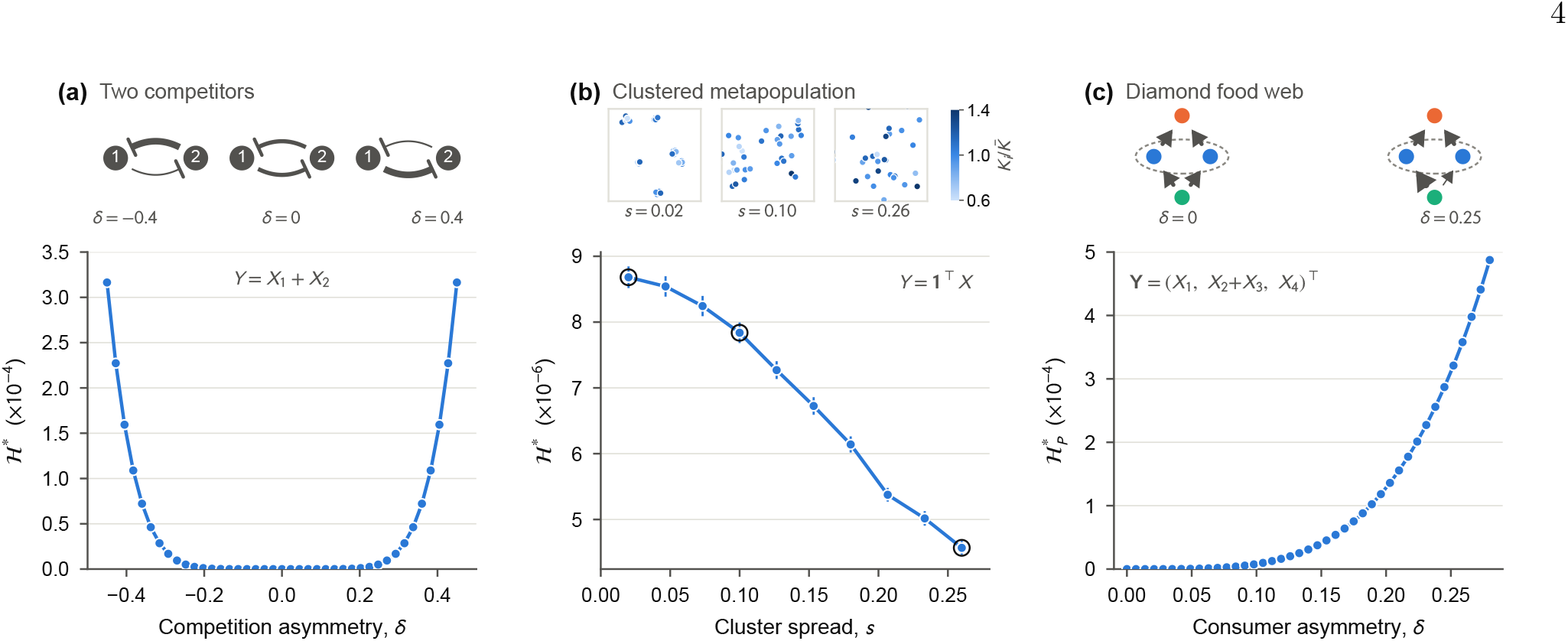
Ecological examples of dynamical heterogeneity; schematics illustrate each system, with line widths indicating interaction strengths. (a) Two-species competition: ℋ* versus interaction asymmetry *δ*. (b) Spatial metapopulation: ℋ* versus cluster spread *s*. Points and error bars show means and standard errors across landscape realizations; insets show representative patch configurations, colored by relative carrying capacity. (c) Diamond food web: 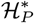 versus consumer asymmetry *δ* for **Y** = (*X*_1_, *X*_2_ + *X*_3_, *X*_4_)^*⊤*^.

For two competing species with self-regulation *A*_11_ = *A*_22_ = *−*1 and asymmetric interspecific terms *A*_12_ = *−*1*/*2 + *δ, A*_21_ = *−*1*/*2 − *δ*, the column sums of *A* are equal only at *δ* = 0, and ℋ* grows symmetrically with |*δ*| [Fig. 2(a)]. In a logistic metapopulation, we aggregate abundance fluctuations across patches with different carrying capacities and distance-dependent dispersal. Smaller cluster spread *s* brings patches within each cluster closer together on average, strengthening their dispersal coupling. The coupled patch dynamics can contribute several relaxation times to regional fluctuations, whereas the scalar OU benchmark has only one. For the parameters considered, we find larger mean ℋ* at smaller *s*, indicating greater spectral mismatch between the regional fluctuation spectrum and its optimal OU approximation [Fig. 2(b)]. Coarse-graining a four-species diamond food web, ordered as a basal resource, two consumers, and a top predator, into three trophic compartments, **Y** = (*X*_1_, *X*_2_ + *X*_3_, *X*_4_)^*⊤*^, extends the comparison to a vector aggregate and its optimal three-dimensional OU model. Here *δ* gives equal and opposite changes in the two resource–consumer interaction strengths: 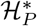 vanishes when the two consumers are dynamically interchangeable, *δ* = 0, and increases with consumer asymmetry [Fig. 2(c)].

### Random community matrices

The previous examples illustrate concrete ecological settings. Random community ensembles reveal how interaction structure affects whether this heterogeneity averages out as richness increases. Following May’s notation, *S* denotes species number in this section, while *S*_*Y*_ (*ω*) remains the aggregate spectrum. Numerical results include only Hurwitz-stable realizations.

In May’s independent-entry ensemble [19], each offdiagonal entry is present with probability *C* and, when present, is drawn independently from *N* (0, *σ*^2^), where *σ* (unrelated to the OU noise amplitude) sets the interaction scale; diagonal entries are *− d*, with common selfregulation rate *d >* 0. For large *S* at fixed connectance *C >* 0, the eigenvalues fill a circular cloud of radius

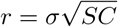

centered at *−d*, so the asymptotic stability boundary is *r* = *d*. For small dimensionless collective interaction strength *r/d*, let 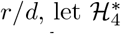 denote the leading quartic term in total-biomass heterogeneity. Its ensemble average satisfies

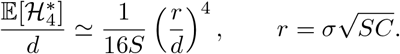

Equivalently, 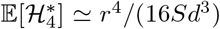 [Fig. 3(a)]. Thus the collective interaction scale *r* sets both the distance to May’s stability boundary and the magnitude of dynamical heterogeneity. The two are not equivalent, however: at fixed *r*, unstructured interaction effects become progressively less visible in total biomass as richness grows, with 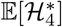 decaying on average as 1*/S* [Fig. 3(c)]. At fixed *r* and *C*, this comparison entails a decrease in typical pairwise interaction strength, 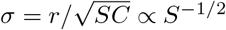.

**FIG. 3.**
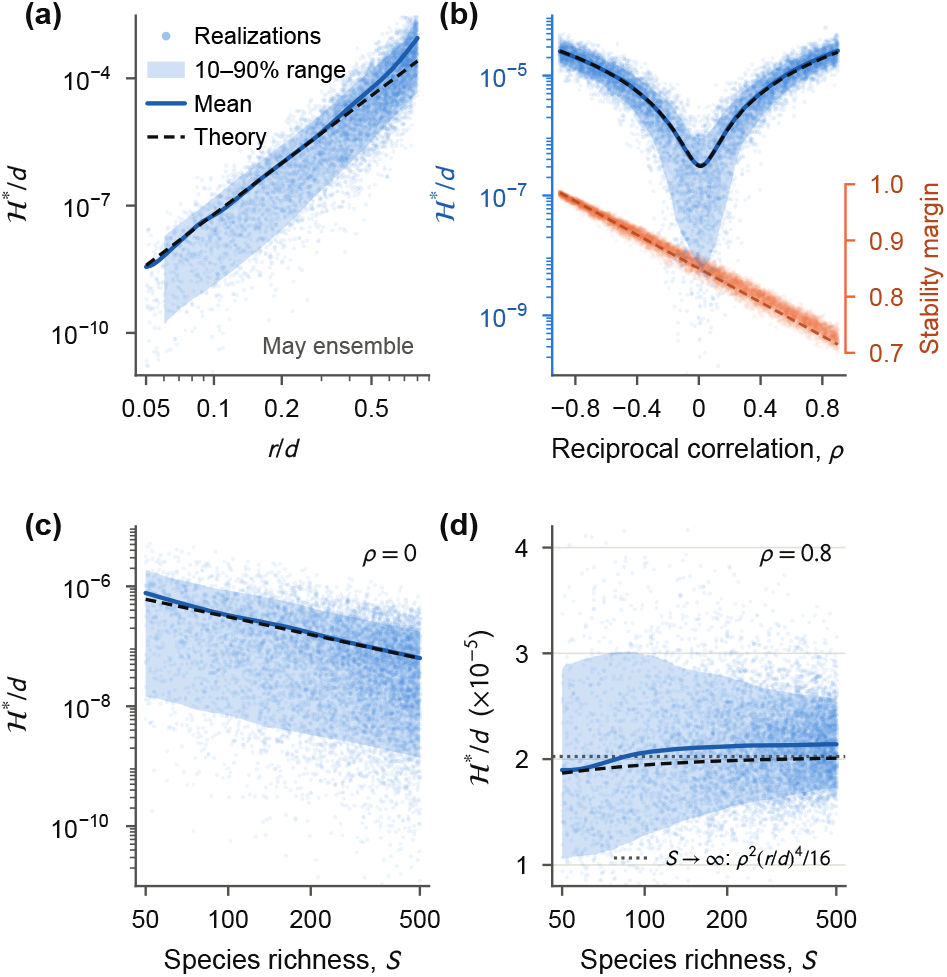
Dynamical heterogeneity and the local-stability margin in random community matrices (*d* = 1, Σ = *I*). Points are individual stable realizations; solid curves and bands show the simulated mean and 10–90% range, and dashed curves the analytical predictions. (a) May ensemble versus *r/d*, with *S* = 100, *C* = 0.3, and 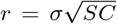. (b) Elliptic ensemble versus reciprocal correlation *ρ*, with *S* = 100 and *r/d* = 0.15; the right axis shows the local-stability margin *−* max Re *λ*(*A*). (c,d) Richness sweeps at *r/d* = 0.15 and 50 *≤S ≤*500, with (c) *ρ* = 0 and (d) *ρ* = 0.8.

Ecological interaction structure changes this relationship. In the fully connected elliptic Gaussian ensemble [20, 21], reciprocal entries have correlation *ρ* = Corr(*A*_*ij*_, *A*_*ji*_). With 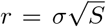, the large-*S* eigenvalue cloud has real and imaginary semi-axes, respectively,

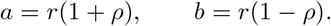

The spectral abscissa is approximately

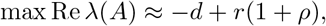

so negative reciprocal, predator–prey-like correlations (*ρ <* 0) increase the local-stability margin by shrinking the horizontal axis of the eigenvalue cloud [21].

Irreducible dynamical heterogeneity responds differently. For small dimensionless collective interaction strength *r/d*, the leading *O*((*r/d*)^4^) prediction, including the first finite-size correction, is

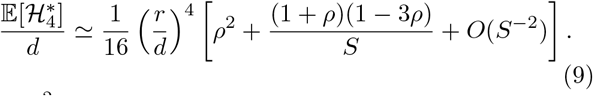

The *ρ*^2^ term is a richness-independent contribution from reciprocal structure, whereas the 1*/S* term gives the first richness-dependent correction. At *ρ* = 0, it reduces to the unstructured result (*r/d*)^4^*/*(16*S*). Within the leading weak-interaction, large-*S* expansion, this correction changes sign at *ρ* = 1*/*3; hence, for sufficiently large *S*, the leading mean heterogeneity decreases with richness for *−* 1 *< ρ <* 1*/*3 and increases for *ρ >* 1*/*3.

For common self-regulation and isotropic forcing, the leading weak-interaction contribution to total-biomass heterogeneity is set by the squared covariance across species between the column sums and row sums of *A*, which describe species’ net effects on total biomass and their responses to a uniform community displacement, respectively [18]. Reciprocity correlates these effects and responses, leaving a contribution that persists as richness grows. Equal positive and negative correlations therefore give the same leading large-*S* heterogeneity, despite opposite effects on stability [Fig. 3(b)].

At fixed *r* and *ρ*, the leading elliptic-law prediction max Re *λ*(*A*) *≈ − d* +*r*(1+*ρ*) is independent of *S*, whereas the effect of richness on aggregate closure depends on reciprocal structure. At *ρ* = 0, reciprocal interactions are uncorrelated, and increasing richness averages away dynamical heterogeneity through the 1*/S* contribution [Fig. 3(c)]. Under sufficiently strong positive reciprocity, *ρ >* 1*/*3, increasing richness reduces the finite-size cancellation of an existing structured contribution, so the predicted heterogeneity increases. The case *ρ* = 0.8 illustrates this reversal [Fig. 3(d)]. Higher-order terms place the simulated mean slightly above this prediction without changing the sign of the richness effect [18]. Thus richness can suppress unstructured contributions or strengthen the dynamical expression of existing interaction structure, while reciprocal correlation itself remains fixed.

Recent microbial work reports that coarse compositional descriptions can become more predictive of community properties as richness increases [22]. That work concerns community-property prediction, with a proposed mechanism distinct from simple averaging. Here, sufficiently strong positive reciprocity reverses the richness effect on temporal OU closure.

## Discussion

Projection methods yield exact reduced equations with memory and colored noise [23–25]. A general coarse-graining isolates the hidden space ker(*P*), whose isolated projected propagator need not decay even when *A* is Hurwitz [18]. Rather than approximate such a memory kernel, we quantify failure of same-dimensional OU closure directly from the aggregate spectrum.

The measure separates structural from dynamical heterogeneity. Composition, spatial structure, and stage structure all shape ecosystem properties such as biomass and variability [26–28], but they obstruct closure only insofar as they remain expressed in the aggregate’s future. A community can be structurally heterogeneous yet dynamically homogeneous with respect to total biomass; conversely, interaction structure alone can force heterogeneity to persist, and neither case is determined by the spectral abscissa that governs asymptotic local stability. An OU or AR(1) fit to total biomass may therefore exhibit systematic spectral lack of fit even in a locally stable community; local stability alone does not justify treating the aggregate as a closed state [4, 6]. A positive ℋ* means that the selected aggregate alone is not a sufficient same-dimensional OU state; it does not exclude an augmented Markov state or a history-dependent reduced model.

The construction is local, applying to stationary fluctuations near a stable equilibrium; strongly state-dependent or nonlinear noise beyond its local additive approximation, temporally correlated forcing, large-amplitude excursions outside the linear neighborhood, limit cycles, regime shifts, and nonlinear memory-driven phenomena lie outside it [29, 30]. That boundary is what makes ℋ* precise rather than general. Within this scope, ℋ* quantifies irreducible dynamical heterogeneity and can also be viewed as a measure of dynamical complexity at the chosen level of aggregation.

## Acknowledgments

We thank Chengyi Long and members of the Savage and Song labs for helpful discussions and insightful feedback. Anthropic’s Claude (opus 5.5) was used to translate the simulation code for Figs. 2 and 3 from MATLAB to Python; the authors verified the translated code against the original MATLAB implementation and the analytical predictions, and take full responsibility for the manuscript.

## Author contributions

M.L. and C.S. designed the study. M.L. developed the theory, performed the analyses and wrote the first draft. V.S. and C.S. revised the manuscript.

## Data availability

The data supporting the findings of this study were generated by numerical simulations. The simulation code and generated data will be deposited in a public repository upon acceptance of the manuscript.

## Appendix A Optimal Ornstein–Uhlenbeck approximation

Let Γ be the stationary community covariance and set *G* = *P* Γ*P* ^*⊤*^, *H* = *PA*Γ*P* ^*⊤*^, and *D*_0_ = *P* Σ*P* ^*⊤*^.

For *τ≥*0, the aggregate covariance *C*_*Y*_ (*τ*) = *Pe*^*Aτ*^ Γ*P* ^*⊤*^ has right derivative *H* at zero. The projected Lyapunov equation gives *H* + *H*^*⊤*^ = *−D*_0_, so 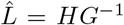 is Hurwitz because 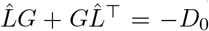, with *G, D*_0_ *≻* 0. The high-frequency tail *S*_*Y*_ (*ω*) *~ D*_0_*/ω*^2^ fixes the OU diffusion at *D*_0_: any different diffusion leaves a positive limiting divergence integrand and hence infinite loss.

For a Hurwitz drift *L*, the OU spectrum is

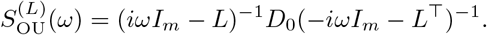

Both spectral matrices are Hermitian: their diagonal entries are real, and the cross-spectra obtained by exchanging two variables are complex conjugates. They are also positive definite under our assumptions. Their generalized eigenvalues *µ*_*j*_(*ω*; *L*) are the values for which a nonzero vector *v*_*j*_ satisfies

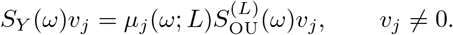

These values are real and positive and generalize the scalar spectral ratio *S*_*Y*_ */S*^(*L*)^. The matrix Itakura–Saito loss [31] is

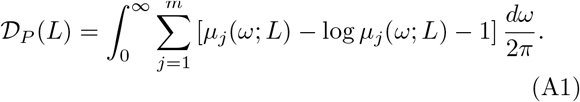

Equivalently, the loss integrand is tr*R −* log det *R − m*, where 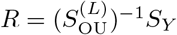.

With diffusion fixed, insert 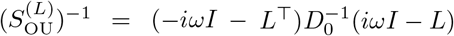 into the spectral loss. Expanding the trace and integrating the covariance and log-determinant terms cancels terms linear in 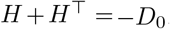, using *H* + *H*^*⊤*^ = *−D*_0_. Thus, for every Hurwitz *L*,

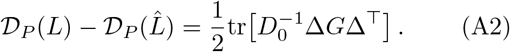

Since *G, D*_0_ *≻* 0, the excess is positive unless Δ = 0, establishing the unique global optimum for both scalar and vector aggregates. The spectral integration details are given in the Supplemental Material.

For a scalar aggregate, *G* = *γ*_*Y*_ (0), 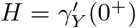, and 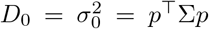. The derivative is taken from the right because white-noise forcing gives the autocovariance a cusp at zero. The projected Lyapunov identity gives 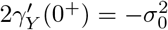, hence

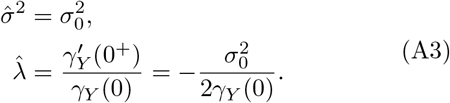

The fit therefore matches the aggregate variance and initial autocorrelation decay rate.

## Appendix B Vanishing heterogeneity and perfect aggregation

If *PAQ* = 0, then *PA* = *LP* with *L* = *PAP* ^+^. Since *PP* ^+^ = *I*_*m*_, *e*^*Lt*^ = *Pe*^*At*^*P* ^+^ decays exponentially, so *L* is Hurwitz. Projecting the full stochastic equation gives 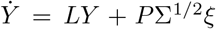, a closed OU process. Thus 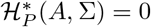 for every Σ *≻* 0.

Conversely, suppose 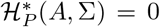 for every Σ *≻* 0. For each forcing, let Γ be its stationary covariance, *G* = *P* Γ*P* ^*⊤*^, and *N* = Γ*P* ^*⊤*^*G*^*−*1^, so *PN* = *I*_*m*_. Equality of spectra gives equality of autocovariance functions, yielding *Pe*^*Aτ*^ *N* = *e*^*Lτ*^ for *τ ≥* 0, where *L* is the matching OU drift. Differentiating once and twice at 0^+^ gives *PAN* = *L* and *PA*^2^*N* = *L*^2^, hence

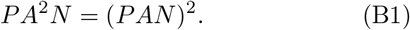

For any hidden direction *q ∈* ker(*P*), set *u* = *PAq* and *V* = *qu*^*⊤*^. Consider the symmetric covariance perturbation

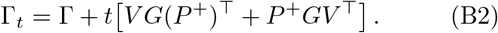

For sufficiently small |*t*|, both Γ_*t*_ and its forcing Σ_*t*_ = *−* (*A*Γ_*t*_ +Γ_*t*_*A*^*⊤*^) remain positive definite. Because *PV* = 0, the aggregate covariance stays *G* and the corresponding right inverse is *N*_*t*_ = *N* + *tV*. The hypothesis therefore requires Eq. (B1) for every such *t*. Its left side is linear in *t*, whereas the quadratic coefficient on the right is (*PAV*)^2^ = (*uu*^*⊤*^)^2^. This must vanish, forcing *u* = 0. Thus *PA* vanishes on every hidden direction, proving *PAQ* = 0.

## References

[1] R.V. O’Neill and B. Rust, Aggregation error in ecological models, Ecological Modelling 7, 91 (1979).

[2] W. G. Cale and P. L. Odell, Behavior of aggregate state variables in ecosystem models, Mathematical Biosciences 49, 121 (1980).

[3] R. H. Gardner, W. G. Cale, and R. V. O’Neill, Robust analysis of aggregation error, Ecology 63, 1771 (1982).

[4] B. Dennis, J. M. Ponciano, S. R. Lele, M. L. Taper, and D. F. Staples, Estimating density dependence, process noise, and observation error, Ecological Monographs 76, 323 (2006).

[5] R. Lande, S. Engen, and B.-E. Sæther, Stochastic Population Dynamics in Ecology and Conservation (Oxford University Press, 2003).

[6] M. Auger-Méthé, K. Newman, D. Cole, F. Empacher, R. Gryba, A. A. King, V. Leos-Barajas, et al., A guide to state-space modeling of ecological time series, Ecological Monographs 91, e01470 (2021).

[7] Y. Iwasa, V. Andreasen, and S. A. Levin, Aggregation in model ecosystems. I. Perfect aggregation, Ecol. Modelling 37, 287 (1987).

[8] H. A. Simon and A. Ando, Aggregation of variables in dynamic systems, Econometrica 29, 111 (1961).

[9] M. Aoki, Control of large-scale dynamic systems by aggregation, IEEE Transactions on Automatic Control 13, 246 (1968).

[10] Y. Iwasa, S. A. Levin, and V. Andreasen, Aggregation in model ecosystems. II. Approximate aggregation, IMA J. Math. Appl. Med. Biol. 6, 1 (1989).

[11] P. Auger and J.-C. Poggiale, Aggregation and emergence in systems of ordinary differential equations, Mathematical and Computer Modelling 27, 1 (1998).

[12] P. Auger, R. Bravo de la Parra, J.-C. Poggiale, E. Sánchez, and L. Sanz, Aggregation methods in dynamical systems and applications in population and community dynamics, Physics of Life Reviews 5, 79 (2008).

[13] L. Barnett and A. K. Seth, Dynamical independence: Discovering emergent macroscopic processes in complex dynamical systems, Phys. Rev. E 108, 014304 (2023).

[14] F. Itakura and S. Saito, Analysis synthesis telephony based on the maximum likelihood method, in Proceedings of the 6th International Congress on Acoustics (1968) pp. C17–C20.

[15] P. Whittle, The analysis of multiple stationary time series, Journal of the Royal Statistical Society. Series B 15, 125 (1953).

[16] D. Kazakos and P. Papantoni-Kazakos, Spectral distance measures between Gaussian processes, IEEE Transactions on Automatic Control 25, 950 (1980).

[17] D. Kazakos, Spectral distance measures between continuous-time vector Gaussian processes, IEEE Transactions on Information Theory 28, 679 (1982).

[18] See Supplemental Material at [URL will be inserted by publisher] for the spectral framework details, extension to general coarse-grainings, discrete-time extension, leading-order scaling derivation for random community matrices, and full model definitions for all numerical examples.

[19] R. M. May, Will a large complex system be stable?, Nature 238, 413 (1972).

[20] H.-J. Sommers, A. Crisanti, H. Sompolinsky, and Y. Stein, Spectrum of large random asymmetric matrices, Phys. Rev. Lett. 60, 1895 (1988).

[21] S. Allesina and S. Tang, Stability criteria for complex ecosystems, Nature 483, 205 (2012).

[22] J. Moran, L. C. Graham, and M. Tikhonov, Emergent predictability in microbial ecosystems, Science 392, eadr1440 (2026).

[23] H. Mori, Transport, collective motion, and Brownian motion, Prog. Theor. Phys. 33, 423 (1965).

[24] R. Zwanzig, Memory effects in irreversible thermodynamics, Phys. Rev. 124, 983 (1961).

[25] A. J. Chorin, O. H. Hald, and R. Kupferman, Optimal prediction and the Mori–Zwanzig representation of irreversible processes, Proceedings of the National Academy of Sciences of the United States of America 97, 2968 (2000).

[26] D. Tilman, The ecological consequences of changes in biodiversity: a search for general principles, Ecology 80, 1455 (1999).

[27] M. Loreau and C. de Mazancourt, Biodiversity and ecosystem stability: a synthesis of underlying mechanisms, Ecology Letters 16, 106 (2013).

[28] S. Wang and M. Loreau, Ecosystem stability in space: α, β and γ variability, Ecology Letters 17, 891 (2014).

[29] M. Scheffer, Critical Transitions in Nature and Society (Princeton University Press, 2009).

[30] F. Ferraro, C. Grilletta, E. Pigani, S. Suweis, S. Azaele, and A. Maritan, Synchronization and chaos in complex ecological communities with delayed interactions (2025), arXiv:2503.21551 [q-bio.PE].

[31] A. Ferrante, C. Masiero, and M. Pavon, Time and spectral domain relative entropy: A new approach to multivariate spectral estimation, IEEE Transactions on Automatic Control 57, 2561 (2012).

