## Supplemental Note for "When does heterogeneity average out in coarse-grained ecological communities?"

### SI. SPECTRAL CONSTRUCTION FOR VECTOR AGGREGATES

We derive the optimal Ornstein–Uhlenbeck (OU) benchmark, the perfect-aggregation criterion, the properties of total-biomass heterogeneity, and the random-matrix predictions, and give the sampled-data benchmark and numerical methods.

Throughout, the full linear stochastic system is

$$\dot{X}(t) = AX(t) + \Sigma^{1/2}\xi(t), \tag{S1}$$

where $X \in \mathbb{R}^n$ denotes deviations about a stable equilibrium, $A \in \mathbb{R}^{n\times n}$ is the community Jacobian and is Hurwitz stable (all eigenvalues have negative real parts), $\Sigma \succ 0$ is the forcing covariance, and $\xi$ is standard Gaussian white noise with $\mathbb{E}[\xi(t)\xi(t')^\top] = I_n\delta(t-t')$. The stationary covariance $\Gamma$ is the unique positive-definite solution of

$$A\Gamma + \Gamma A^\top + \Sigma = 0. \tag{S2}$$

For a coarse-graining $Y = PX \in \mathbb{R}^m$, with aggregation matrix $P \in \mathbb{R}^{m\times n}$ of full row rank, the exact stationary cross-spectrum is

$$S_Y(\omega) = P(i\omega I_n - A)^{-1}\Sigma(-i\omega I_n - A^\top)^{-1}P^\top. \tag{S3}$$

We use the usual two-sided spectral density. For a real vector stationary process, the cross-spectral matrix satisfies

$$S_Y(-\omega) = S_Y(\omega)^\top = \overline{S_Y(\omega)}. \tag{S4}$$

Thus a vector cross-spectrum need not itself be an even matrix-valued function. The scalar matrix-divergence integrand defined below is nevertheless even because trace and determinant are invariant under transposition. We may therefore evaluate the divergence over $\omega \geq 0$, where $\omega$ is angular frequency. For a scalar aggregate, let $\gamma_Y(\tau) = \mathbb{E}[Y(t+\tau)Y(t)]$ be its stationary autocovariance. Then $S_Y(-\omega) = S_Y(\omega)$, and the stationary variance is

$$\gamma_Y(0) = \int_{-\infty}^{\infty} S_Y(\omega)\frac{\mathrm{d}\omega}{2\pi} = 2\int_0^{\infty} S_Y(\omega)\frac{\mathrm{d}\omega}{2\pi}. \tag{S5}$$

#### SI.1. Matrix Itakura–Saito divergence

The same-dimensional linear-Gaussian benchmark for a vector aggregate is an $m$-dimensional Ornstein–Uhlenbeck process

$$\dot{Y} = LY + \zeta(t), \qquad \langle\zeta(t)\zeta(t')^\top\rangle = D\,\delta(t-t'), \tag{S6}$$

with Hurwitz drift $L$, diffusion matrix $D \succ 0$, and zero-mean Gaussian white noise $\zeta$. Its spectrum is

$$S_{\mathrm{OU}}^{(L,D)}(\omega) = (i\omega I_m - L)^{-1} D (-i\omega I_m - L^\top)^{-1}. \tag{S7}$$

For Hermitian positive-definite spectral matrices, the matrix Itakura–Saito divergence [1, 2] is

$$\mathcal{D}_P(L, D) = \int_0^\infty \Big[ \mathrm{tr}\Big( S_{\mathrm{OU}}^{(L,D)}(\omega)^{-1} S_Y(\omega) \Big) - \log\det\Big( S_{\mathrm{OU}}^{(L,D)}(\omega)^{-1} S_Y(\omega) \Big) - m \Big] \frac{\mathrm{d}\omega}{2\pi}. \tag{S8}$$

Equivalently, if $\mu_1(\omega), \ldots, \mu_m(\omega)$ are the generalized eigenvalues of the Hermitian pair $(S_Y(\omega), S_{\mathrm{OU}}^{(L,D)}(\omega))$, the integrand is

$$\sum_{j=1}^{m} \left[ \mu_j(\omega) - \log \mu_j(\omega) - 1 \right], \tag{S9}$$

which is nonnegative and vanishes only when the two spectral matrices agree at that frequency.

The high-frequency expansion of Eq. (S3) is

$$S_Y(\omega) = \frac{P \Sigma P^\top}{\omega^2} + O(\omega^{-3}), \qquad \omega \to \infty. \tag{S10}$$

Similarly, $S_{\mathrm{OU}}^{(L,D)}(\omega) = D/\omega^2 + O(\omega^{-3})$. If $D \neq P\Sigma P^\top$, the generalized eigenvalues in Eq. (S9) tend to constants that are not all equal to one, so the integrand tends to a positive constant and the divergence is infinite. Therefore any finite-divergence OU benchmark must have

$$D_0 = P \Sigma P^\top. \tag{S11}$$

With this tail matched, $\mu_j(\omega) = 1 + O(\omega^{-1})$, so $\mu_j - \log \mu_j - 1 = O(\omega^{-2})$ and the divergence is finite.

### SI.2. Optimal OU drift

Define the aggregate covariance $G$ and the covariance derivative $H$ by

$$G = P \Gamma P^\top, \qquad H = P A \Gamma P^\top. \tag{S12}$$

Multiplying Eq. (S2) by $P$ and $P^\top$ gives

$$H + H^\top + D_0 = 0. \tag{S13}$$

The right derivative at zero of the exact aggregate covariance $C_Y(\tau) = \mathbb{E}[Y(t+\tau) Y(t)^\top]$ is $H$. Define

$$\hat{L} = H G^{-1} = P A \Gamma P^\top (P \Gamma P^\top)^{-1}. \tag{S14}$$

It satisfies

$$\hat{L} G + G \hat{L}^\top + D_0 = 0. \tag{S15}$$

Since $G \succ 0$ and $D_0 \succ 0$, the Lyapunov theorem implies that $\hat{L}$ is Hurwitz.

**Lemma 1** (Optimal vector OU drift)**.** *For every Hurwitz $L$,*

$$\mathcal{D}_P(L, D_0) - \mathcal{D}_P(\hat{L}, D_0) = \frac{1}{2}\mathrm{tr}\Big[D_0^{-1}(L - \hat{L})G(L - \hat{L})^\top\Big]. \tag{S16}$$

*Consequently, $\hat{L}$ is the unique minimizer of the matrix Itakura–Saito divergence among stable OU drifts with diffusion $D_0$.*

*Proof.* Set $K = \hat{L}$, $\Delta = L - K$, and

$$M = \frac{1}{2}(H - H^\top).$$

The required spectral moments are

$$\begin{aligned} \int_{-\infty}^{\infty} S_Y(\omega)\frac{\mathrm{d}\omega}{2\pi} &= G, \\ \mathrm{p.\,v.}\int_{-\infty}^{\infty} i\omega S_Y(\omega)\frac{\mathrm{d}\omega}{2\pi} &= M. \end{aligned} \tag{S17}$$

Here p. v. denotes the Cauchy principal value. The second identity is the symmetric Fourier value at the cusp of the covariance function: its right and left derivatives at zero are $H$ and $-H^\top$, respectively.

For real Hurwitz matrices $L$ and $K$, factorization over their eigenvalues gives

$$\int_0^{\infty} \log \frac{|\det(i\omega I_m - L)|^2}{|\det(i\omega I_m - K)|^2}\frac{\mathrm{d}\omega}{2\pi} = -\frac{1}{2}\mathrm{tr}(L - K). \tag{S18}$$

Using a symmetric frequency cutoff, expanding

$$S_{\mathrm{OU}}^{(L,D_0)}(\omega)^{-1} = (-i\omega I_m - L^\top)D_0^{-1}(i\omega I_m - L),$$

and then applying Eqs. (S17) and (S18) yields

$$\begin{aligned} \mathcal{D}_P(L, D_0) - \mathcal{D}_P(K, D_0) = \mathrm{tr}(D_0^{-1}\Delta M) \\ + \frac{1}{2}\mathrm{tr}\Big[D_0^{-1}\left(LGL^\top - KGK^\top\right)\Big] + \frac{1}{2}\mathrm{tr}(\Delta). \end{aligned} \tag{S19}$$

Because $KG = H$, expanding $L = K + \Delta$ shows that the terms linear in $\Delta$ are

$$\mathrm{tr}\Big[D_0^{-1}\Delta(M + H^\top)\Big] + \frac{1}{2}\mathrm{tr}(\Delta).$$

Equation (S13) implies

$$M + H^\top = \frac{1}{2}(H + H^\top) = -\frac{1}{2}D_0,$$

so these linear terms cancel. The remaining term is

$$\frac{1}{2}\mathrm{tr}\Big[D_0^{-1}\Delta G\Delta^\top\Big],$$

which is Eq. (S16). It is nonnegative and vanishes only when $\Delta = 0$. □

Combining the high-frequency condition $D = D_0 = P\Sigma P^\top$ with Lemma 1, the vector irreducible dynamical heterogeneity is

$$\mathcal{H}_P^*(A, \Sigma) = \min_{\substack{L \text{ Hurwitz} \\ D \succ 0}} \mathcal{D}_P(L, D) = \mathcal{D}_P(\hat{L}, D_0). \tag{S20}$$

#### SI.3. Scalar specialization

For $P = p^\top$, $p \neq 0$, write $\mathcal{D}(\lambda, \sigma^2) = \mathcal{D}_{p^\top}(\lambda, \sigma^2)$ and $\sigma_0^2 = p^\top \Sigma p$. The autocovariance $\gamma_Y(\tau) = p^\top e^{A\tau}\Gamma p$ for $\tau \geq 0$ satisfies $2\gamma_Y'(0^+) + \sigma_0^2 = 0$, so Eq. (S14) becomes

$$\hat{\lambda} = -\frac{\sigma_0^2}{2\gamma_Y(0)} = \frac{\gamma_Y'(0^+)}{\gamma_Y(0)}. \tag{S21}$$

Define $\mathcal{H}_p^* = \mathcal{D}(\hat{\lambda}, \sigma_0^2)$; without a subscript, $\mathcal{H}^* = \mathcal{H}_{\mathbf{1}}^*$ denotes the heterogeneity of total biomass, with $\mathbf{1}$ the all-ones vector. Lemma 1 directly gives, for every $\lambda < 0$,

$$\mathcal{D}(\lambda, \sigma_0^2) - \mathcal{H}_p^* = \frac{\gamma_Y(0)}{2\sigma_0^2}(\lambda - \hat{\lambda})^2. \tag{S22}$$

For one forcing covariance, $\mathcal{H}_P^* = 0$ identifies an OU stationary marginal law, which need not imply deterministic closure; the structural criterion requires all positive-definite forcing covariances.

### SII. VANISHING HETEROGENEITY AND PERFECT AGGREGATION

The condition $PAQ = 0$, equivalently $PA = LP$ for some $L$, is the perfect-aggregation condition of Iwasa, Andreasen, and Levin [3].

**Theorem 1.** *Let $A \in \mathbb{R}^{n\times n}$ be Hurwitz stable and $P \in \mathbb{R}^{m\times n}$ have full row rank. With $P^+$ the Moore–Penrose pseudoinverse and $Q = I_n - P^+P$ the orthogonal projection onto* $\ker(P)$*,*

$$\mathcal{H}_P^*(A, \Sigma) = 0 \text{ for every } \Sigma \succ 0 \iff PAQ = 0. \tag{S23}$$

*Proof.* If $PAQ = 0$, then $PA = LP$ with $L = PAP^+$. Since $PP^+ = I_m$, $e^{Lt} = Pe^{At}P^+$ decays exponentially, so $L$ is Hurwitz. Projecting Eq. (S1) gives $\dot{Y} = LY + P\Sigma^{1/2}\xi$, a closed OU process with diffusion $P\Sigma P^\top$. Hence $\mathcal{H}_P^*(A, \Sigma) = 0$ for every $\Sigma \succ 0$.

Conversely, let $\mathcal{H}_P^*(A, \Sigma) = 0$ for every $\Sigma \succ 0$. For any such forcing, let $\Gamma$ solve Eq. (S2), set $G = P\Gamma P^\top$, and define $N = \Gamma P^\top G^{-1}$, so $PN = I_m$. Zero divergence implies equality of the continuous aggregate and OU spectra and therefore of their covariance functions for some Hurwitz drift $L$:

$$Pe^{A\tau}N = e^{L\tau}, \qquad \tau \geq 0. \tag{S24}$$

The first two right derivatives at zero give $PAN = L$ and $PA^2N = L^2$, hence

$$PA^2N = (PAN)^2. \tag{S25}$$

Fix $q \in \ker(P)$, set $u = PAq$, and let $V = qu^\top$, so $PV = 0$. Vary the stationary covariance as

$$\Gamma_t = \Gamma + t\left[VG(P^+)^\top + P^+GV^\top\right]. \tag{S26}$$

For sufficiently small $|t|$, both $\Gamma_t$ and $\Sigma_t = -(A\Gamma_t + \Gamma_t A^\top)$ remain positive definite by continuity. Because $PV = 0$ and $PP^+ = I_m$,

$$P\Gamma_t P^\top = G, \qquad N_t = \Gamma_t P^\top G^{-1} = N + tV. \tag{S27}$$

The all-forcing hypothesis requires Eq. (S25) for every such $t$. Its left side is linear in $t$, while the quadratic coefficient on the right is

$$(PAV)^2 = (uu^\top)^2 = (u^\top u)uu^\top. \tag{S28}$$

This coefficient must vanish, so $u = PAq = 0$. Since $q$ was any hidden direction, $PAQ = 0$. □

**Remark 1** (Why one forcing covariance is insufficient)**.** The quantifier over forcing covariances cannot be dropped, even when the single covariance is isotropic. Let

$$A = \begin{pmatrix} -1 & -4 \\ 0 & -3 \end{pmatrix}, \qquad P = \begin{pmatrix} 1 & 1 \end{pmatrix}, \qquad \Sigma = I_2.$$

Perfect aggregation fails because

$$PA = (-1, -7)$$

is not proportional to $P$. Nevertheless,

$$P(sI_2 - A)^{-1} = \left( \frac{1}{s+1}, \frac{s-3}{(s+1)(s+3)} \right),$$

and hence

$$\begin{aligned} S_Y(\omega) &= \frac{1}{\omega^2+1} + \frac{\omega^2+9}{(\omega^2+1)(\omega^2+9)} \\ &= \frac{2}{\omega^2+1}. \end{aligned} \tag{S29}$$

Thus the stationary marginal process $Y$ has the same law as an OU process with rate $-1$ and diffusion intensity $P\Sigma P^\top = 2$, so $\mathcal{H}^*_P(A, I_2) = 0$ despite the failure of perfect aggregation. The hidden mode survives in the transfer function through the all-pass factor $(s-3)/(s+3)$, whose modulus is one on the imaginary axis, and is therefore invisible in the aggregate power spectrum.

### SIII. PROPERTIES OF TOTAL-BIOMASS HETEROGENEITY

We consider the stationary system in Eq. (S1) with $n$ species, a Hurwitz-stable community matrix $A$, and independent environmental forcing of unit intensity on each species. The aggregate is total biomass, $Y(t) = \mathbf{1}^\top X(t)$, where $\mathbf{1} \in \mathbb{R}^n$ is the all-ones vector. Let $S_Y(\omega)$ be its two-sided power spectrum at angular frequency $\omega$, and let $\gamma_Y(\tau) = \mathbb{E}[Y(t+\tau)Y(t)]$ be its autocovariance at time lag $\tau$. Define the normalized spectrum $q$ and normalized variance $v > 0$ by

$$q(\omega) = \frac{S_Y(\omega)}{n}, \qquad v = \frac{\gamma_Y(0)}{n} = \int_{-\infty}^{\infty} q(\omega) \frac{d\omega}{2\pi}. \tag{S30}$$

The aggregate diffusion intensity is $n$. Thus the optimal Ornstein–Uhlenbeck (OU) rate and spectrum are

$$\hat{\lambda} = -\frac{1}{2v}, \qquad S_{\mathrm{OU}}(\omega) = \frac{n}{\omega^2 + \hat{\lambda}^2}. \tag{S31}$$

Writing $R(\omega) = S_Y(\omega)/S_{\mathrm{OU}}(\omega)$ for the spectral ratio and $f(u) = u - \log u - 1$ for $u > 0$, the minimized spectral loss is

$$\mathcal{H}^* = \int_0^\infty f(R(\omega)) \frac{d\omega}{2\pi}, \qquad R(\omega) = (\omega^2 + \hat{\lambda}^2) q(\omega). \tag{S32}$$

The following properties follow from this definition.

#### SIII.1. Finiteness, nonnegativity, and the zero case

The scalar spectrum is continuous and strictly positive for real $\omega$. Because it is even, its high-frequency expansion has the form

$$q(\omega) = \omega^{-2} + O(\omega^{-4}), \qquad R(\omega) = 1 + O(\omega^{-2}). \tag{S33}$$

Since $f(1+\epsilon) = \epsilon^2/2 + O(\epsilon^3)$ as $\epsilon \to 0$, the loss integrand is $O(\omega^{-4})$ and is integrable. Also $f(u) \geq 0$, with equality only at $u = 1$, so $0 \leq \mathcal{H}^* < \infty$. Continuity of $R$ implies that $\mathcal{H}^* = 0$ if and only if $S_Y = S_{\mathrm{OU}}$ at every frequency. A centered stationary Gaussian process is determined by its covariance function, so this is equivalent to the aggregate having the same stationary process law as the fitted OU process. It does not imply deterministic closure for this single forcing covariance; the distinction is established in Sec. SII.

#### SIII.2. Normalized spectrum and invariance under relabeling

Equations (S30)–(S32) show that $q$ determines both the fitted rate and $\mathcal{H}^*$. Systems with the same normalized spectrum therefore have the same heterogeneity, even if they have different dimensions. In particular, relabeling species replaces $A$ by $\Pi A \Pi^\top$, where $\Pi$ is a permutation matrix. Since $\Pi^\top \mathbf{1} = \mathbf{1}$ and the forcing covariance is unchanged, the aggregate spectrum, and hence $\mathcal{H}^*$, is unchanged.

#### SIII.3. Continuity within the stable domain

For fixed $n$, $\mathcal{H}^*$ is continuous in $A$ throughout the Hurwitz-stable domain. To see this, take a compact neighborhood of any stable matrix contained in that domain. The stationary covariance $\Gamma$, defined by Eq. (S2), varies continuously there, as do $v > 0$ and $\hat{\lambda}$. On bounded frequency intervals, $R$ is jointly continuous in $A$ and $\omega$ and bounded away from zero. The resolvent expansion gives Eq. (S33) uniformly on the chosen neighborhood, so $f(R) = O(\omega^{-4})$ uniformly at high frequencies. Dominated convergence then establishes continuity of the integral in Eq. (S32).

#### SIII.4. Scaling all dynamical rates

Let $c > 0$ multiply the community matrix, $A \mapsto cA$, while the forcing intensity remains fixed. Use a subscript $c$ for quantities in the rescaled system. The resolvent formula and Lyapunov equation give

$$S_{Y,c}(\omega) = c^{-2} S_Y(\omega/c), \qquad \gamma_{Y,c}(0) = c^{-1}\gamma_Y(0), \qquad \hat{\lambda}_c = c\hat{\lambda}. \tag{S34}$$

Consequently, $R_c(\omega) = R(\omega/c)$. Changing variables in the loss integral gives $\mathcal{H}^*_c = c\mathcal{H}^*$, consistent with the inverse-time units of the measure.

#### SIII.5. Invariance under independent replication

Consider $k \geq 2$ isolated copies of the same community, where $k$ is an integer, each with matrix $A$ and independent realizations of the same unit-intensity forcing. Let $Y_j$ be the total biomass of copy $j$, and let $Y_{\rm rep} = \sum_{j=1}^{k} Y_j$ be their combined total biomass. Independence gives

$$S_{\rm rep}(\omega) = kS_Y(\omega), \qquad \gamma_{\rm rep}(0) = k\gamma_Y(0), \tag{S35}$$

where $S_{\rm rep}$ and $\gamma_{\rm rep}$ are the spectrum and autocovariance of $Y_{\rm rep}$. Its diffusion intensity is $kn$, so its fitted rate is

$$\hat{\lambda}_{\rm rep} = -\frac{kn}{2k\gamma_Y(0)} = \hat{\lambda}. \tag{S36}$$

The optimal OU spectrum therefore also increases by the factor $k$. The spectral ratio is unchanged at every frequency, giving $\mathcal{H}^*_{\rm rep} = \mathcal{H}^*$, where $\mathcal{H}^*_{\rm rep}$ is the minimized loss for the combined scalar total. In particular, duplicating a community into two isolated copies leaves its total-biomass heterogeneity unchanged.

### SIV. DISCRETE-TIME FORMULATION AND SAMPLED-DATA BENCHMARK

The same construction has a discrete-time analogue. Consider

$$X_{t+1} = A_d X_t + \varepsilon_t, \tag{S37}$$

where $A_d$ is Schur stable (all eigenvalues have modulus less than one) and the innovations $\varepsilon_t$ are independent, zero-mean Gaussian vectors with covariance $\Sigma_d \succ 0$. For a scalar aggregate $Y_t = p^\top X_t$, $p \neq 0$, the stationary spectrum is

$$S_Y(\omega) = p^\top (e^{i\omega} I - A_d)^{-1} \Sigma_d (e^{-i\omega} I - A_d^\top)^{-1} p, \qquad \omega \in [-\pi, \pi]. \tag{S38}$$

The same-dimensional discrete-time comparator is a first-order autoregressive [AR(1)] process,

$$Y_{t+1} = \phi Y_t + \epsilon_t, \qquad |\phi| < 1, \tag{S39}$$

with independent Gaussian innovations of variance $\sigma_\epsilon^2 > 0$ and spectrum

$$S_{\rm AR}^{(\phi,\sigma_\epsilon^2)}(\omega) = \frac{\sigma_\epsilon^2}{|e^{i\omega} - \phi|^2}. \tag{S40}$$

The discrete-time Itakura–Saito divergence is

$$\mathcal{D}_{\rm d}(\phi, \sigma_\epsilon^2) = \frac{1}{2} \int_{-\pi}^{\pi} \left[ \frac{S_Y(\omega)}{S_{\rm AR}^{(\phi,\sigma_\epsilon^2)}(\omega)} - \log \frac{S_Y(\omega)}{S_{\rm AR}^{(\phi,\sigma_\epsilon^2)}(\omega)} - 1 \right] \frac{{\rm d}\omega}{2\pi}. \tag{S41}$$

The factor $1/2$ makes the full-frequency integral consistent with the positive-frequency convention in Eq. (S8). The discrete-time heterogeneity is

$$\mathcal{H}^*_{\rm d} = \inf_{\substack{|\phi|<1 \\ \sigma_\epsilon^2 > 0}} \mathcal{D}_{\rm d}(\phi, \sigma_\epsilon^2). \tag{S42}$$

For a scalar aggregate, exact aggregation gives an AR(1) process and hence $\mathcal{H}^*_{\rm d} = 0$; the converse need not hold for one specified innovation covariance.

#### SIV.1. Sampled-data benchmark

The sampled-data benchmark in Fig. S1 tests convergence of the OU benchmark estimated from sampled aggregate observations. Let $T$ denote the observation-window duration, $\Delta t > 0$ the sampling interval, and $N_{\mathrm{obs}}$ the number of retained observations. Each trial starts from the stationary distribution $X_0 \sim \mathcal{N}(0, \Gamma)$, so no burn-in is used. The MATLAB random-number generator is initialized with seed 42. We retain $N_{\mathrm{obs}} = \mathrm{round}(T/\Delta t) + 1$ observations, including both endpoints; $T/\Delta t$ is an integer for the grid used here. Writing $X_k = X(k\Delta t)$, the continuous-time full system is discretized exactly:

$$X_{k+1} = e^{A\Delta t} X_k + \eta_k, \tag{S43}$$

where the innovations $\eta_k \sim \mathcal{N}(0, \Gamma_{\Delta t})$ are independent across steps and independent of $X_0$, with

$$\Gamma_{\Delta t} = \int_0^{\Delta t} e^{A\tau} \Sigma e^{A^\top \tau} \, \mathrm{d}\tau. \tag{S44}$$

Only $Y_k = Y(k\Delta t) = p^\top X_k$, for $k = 0, \ldots, N_{\mathrm{obs}} - 1$, is retained. A zero-mean AR(1) model is fitted without demeaning or an intercept by ordinary least squares,

$$\hat{\phi} = \frac{\sum_{k=0}^{N_{\mathrm{obs}}-2} Y_{k+1} Y_k}{\sum_{k=0}^{N_{\mathrm{obs}}-2} Y_k^2}, \tag{S45}$$

then converted to a continuous-time relaxation rate when $0 < \hat{\phi} < 1$:

$$\hat{\lambda}_{\mathrm{OU}} = \frac{\log \hat{\phi}}{\Delta t}. \tag{S46}$$

The population AR(1) coefficient at sampling interval $\Delta t$ is

$$\phi_{\Delta t}^{\mathrm{pop}} = \frac{\gamma_Y(\Delta t)}{\gamma_Y(0)} = \frac{p^\top e^{A\Delta t} \Gamma p}{p^\top \Gamma p}. \tag{S47}$$

When $\phi_{\Delta t}^{\mathrm{pop}} > 0$, the corresponding real continuous-time rate is

$$\lambda_{\Delta t}^{\mathrm{pop}} = \frac{\log \phi_{\Delta t}^{\mathrm{pop}}}{\Delta t}. \tag{S48}$$

As $\Delta t \to 0$, $\lambda_{\Delta t}^{\mathrm{pop}} \to \gamma_Y'(0^+)/\gamma_Y(0)$, which equals the continuous-time OU benchmark rate. At fixed $\Delta t$ with $\phi_{\Delta t}^{\mathrm{pop}} > 0$, the fitted rate converges to $\lambda_{\Delta t}^{\mathrm{pop}}$ as $T \to \infty$; finer sampling at fixed $T$ does not eliminate finite-window estimation uncertainty. For each sampled realization with $0 < \hat{\phi} < 1$, the benchmark lack of fit is

$$\widehat{\mathcal{H}}_{T,\Delta t} = \mathcal{D}(\hat{\lambda}_{\mathrm{OU}}, \sigma_0^2), \tag{S49}$$

while the finite-sampling-interval population target is

$$\mathcal{H}_{\Delta t}^{\mathrm{pop}} = \mathcal{D}(\lambda_{\Delta t}^{\mathrm{pop}}, \sigma_0^2). \tag{S50}$$

Both quantities can equivalently be evaluated using Eq. (S22). They use the continuous-time divergence, not $\mathcal{H}_{\mathrm{d}}^*$. The diffusion is fixed at $\sigma_0^2 = p^\top \Sigma p$, and losses are evaluated against the known

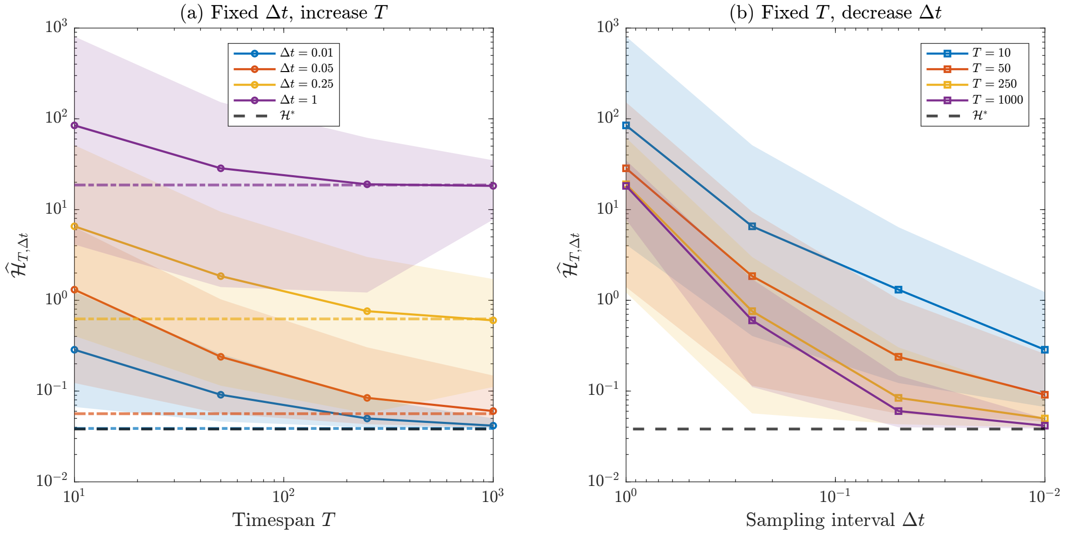


FIG. S1. Sampled-data convergence of the fitted OU benchmark for irreducible dynamical heterogeneity. The OU rate is fitted to sampled aggregate observations; diffusion and the comparison spectrum are known. (a) The sampling interval is fixed and the observation window $T$ increases. (b) The observation window is fixed and $\Delta t$ decreases. Colored dash-dotted lines in panel (a) denote the finite-$\Delta t$ population targets $\mathcal{H}^{\mathrm{pop}}_{\Delta t}$, and the dashed black line denotes the continuous-time value $\mathcal{H}^*$. Solid curves show medians and shaded regions show 10–90% quantiles over retained valid fits.

continuous-time spectrum. This tests convergence of the OU fit; a fully empirical estimate would additionally require estimating $S_Y(\omega)$. Observation error is not included.

The sampled-data benchmark evaluates the continuous-time divergence by trapezoidal quadrature on $N_\omega = 2000$ nodes $x_k = k/N_\omega$, $k = 0, \ldots, N_\omega - 1$, after the transformation $\omega = x/(1-x)$, including its Jacobian $1/(1-x)^2$.

The benchmark uses the two-species matrix

$$A = \begin{pmatrix} -1.1 + \varepsilon & -1 \\ -1 & -1.1 - \varepsilon \end{pmatrix}, \qquad \varepsilon = 0.2, \tag{S51}$$

with $\Sigma = I_2$ and $Y = X_1 + X_2$. Observation windows are $T \in \{10, 50, 250, 1000\}$, and sampling intervals are $\Delta t \in \{0.01, 0.05, 0.25, 1.0\}$. Each combination uses 500 independent attempted trials. Fits with $\hat{\phi} \notin (0, 1)$, nonpositive mean squared residuals, or nonfinite estimates are discarded. Curves show medians and shaded regions show 10–90% quantiles over retained fits. At $\Delta t = 1$, 372 of 500 trials are retained for $T = 10$ and 472 of 500 for $T = 50$; all trials are retained for the other grid points.

### SV. PROJECTED HIDDEN DYNAMICS NEED NOT BE STABLE

The projected dynamics on $\ker(P)$ can be unstable even when the full community is stable. A two-dimensional example is

$$A = \begin{pmatrix} -3 & 10 \\ -1 & 1 \end{pmatrix}, \qquad P = \begin{pmatrix} 1 & 0 \end{pmatrix}, \qquad Q = \begin{pmatrix} 0 & 0 \\ 0 & 1 \end{pmatrix}. \tag{S52}$$

The eigenvalues of $A$ are $-1 \pm i\sqrt{6}$, so the full system is Hurwitz stable. The restriction of $QAQ$ to $\ker(P)$ has eigenvalue $+1$, and its propagator grows as $e^t$. The full resolvent $(i\omega I - A)^{-1}$ therefore avoids assuming stability of the projected hidden dynamics.

### SVI. RANDOM-MATRIX CALCULATION

This section derives the weak-interaction predictions for total-biomass heterogeneity in main-text Fig. 3. With common self-regulation $d > 0$, the leading term $\mathcal{H}_4^*/d$ is quartic in the collective interaction strength $r/d$, with $r$ defined for each ensemble below. Ensemble averages of this quartic term are unconditioned on stability; numerical spectral evaluations retain only Hurwitz-stable realizations (Sec. SVII SVII.1).

#### SVI.1. Resolvent expansion for a fixed interaction matrix

Write

$$A = -dI + \sigma B, \tag{S53}$$

Here $\sigma \geq 0$ scales interaction strengths, $B$ is a dimensionless $S \times S$ matrix with zero diagonal, and $p = \mathbf{1}$. In this section, $S$ denotes species richness, consistent with the main text, and $\Sigma = I_S$. With $g(\omega) = (i\omega + d)^{-1}$, the Neumann expansion is

$$(i\omega I - A)^{-1} = g(\omega) \sum_{k=0}^{\infty} \big(\sigma g(\omega) B\big)^k. \tag{S54}$$

The condition $\sigma \|B\|_2/d < 1$, where $\|B\|_2$ is the Euclidean operator norm, ensures uniform convergence over real $\omega$. Remainder estimates below are for fixed $B$ and $S$ as $\sigma/d \to 0$. For the random ensembles considered below, $\sigma\|B\|_2/d$ is typically of order $r/d$.

The aggregate spectrum expands as

$$S_Y(\omega) = |g|^2 \sum_{q=0}^{\infty} \sigma^q \sum_{k+\ell=q} g^k \bar{g}^\ell \mathbf{1}^\top B^k (B^\top)^\ell \mathbf{1}. \tag{S55}$$

Writing $S_Y(\omega) = \sum_{q=0}^{\infty} \sigma^q S_Y^{(q)}(\omega)$, the zeroth-order term is $S_Y^{(0)}(\omega) = S/(\omega^2 + d^2)$. The first-order coefficient is

$$S_Y^{(1)}(\omega) = |g|^2 (g + \bar{g}) b_{\text{tot}} = \frac{2d\, b_{\text{tot}}}{(\omega^2 + d^2)^2}, \qquad b_{\text{tot}} = \mathbf{1}^\top B \mathbf{1}. \tag{S56}$$

This contribution has the same shape as a first-order shift of the OU relaxation rate and is absorbed by the OU fit.

Let

$$c_j = \sum_i B_{ij}, \qquad u_j = \sum_k B_{jk} \tag{S57}$$

be the column and row sums of $B$, and let

$$\tilde{c}_j = c_j - \bar{c}, \qquad \tilde{u}_j = u_j - \bar{u}, \qquad \bar{c} = \bar{u} = b_{\text{tot}}/S. \tag{S58}$$

With $\langle u, c\rangle = \sum_j u_j c_j$ and $\|c\|^2 = \sum_j c_j^2$, the second-order coefficient is

$$S_Y^{(2)}(\omega) = \frac{2(d^2 - \omega^2)\langle u, c\rangle}{(\omega^2 + d^2)^3} + \frac{\|c\|^2}{(\omega^2 + d^2)^2}. \tag{S59}$$

Define $q_c = \langle \tilde{u}, \tilde{c} \rangle = \langle u, c \rangle - b_{\text{tot}}^2/S$. After fitting the OU rate, the leading relative spectral mismatch is

$$\frac{S_Y(\omega)}{S_{\text{OU}}(\omega)} - 1 = \frac{\sigma^2(d^2 - 3\omega^2)q_c}{S(\omega^2 + d^2)^2} + O(\sigma^3). \tag{S60}$$

Using $x - \log x - 1 = (x-1)^2/2 + O((x-1)^3)$, we obtain

$$\mathcal{H}^* = \frac{\sigma^4 q_c^2}{2S^2} \int_0^\infty \frac{(d^2 - 3\omega^2)^2}{(\omega^2 + d^2)^4} \frac{\mathrm{d}\omega}{2\pi} + O(\sigma^5/d^4). \tag{S61}$$

The integral without the factor $1/(2\pi)$ equals $\pi/(4d^3)$, giving

$$\mathcal{H}_4^* = \frac{\sigma^4 q_c^2}{16 S^2 d^3}, \qquad \mathcal{H}^* = \mathcal{H}_4^* + O(\sigma^5/d^4). \tag{S62}$$

Here $q_c/S$ is the covariance across species of the column and row sums of $B$, normalized by $1/S$. The corresponding sums of $A$ measure species' net effects on total biomass and their responses to a uniform community displacement. The quartic term $\mathcal{H}_4^*$ is therefore proportional to the squared covariance of these effects and responses. A vanishing covariance removes this leading term but does not in general imply perfect aggregation.

#### SVI.2. May's independent-entry ensemble

In May's ensemble [4], each directed off-diagonal entry of $B$ is present with fixed probability $C > 0$ and, when present, is drawn from $\mathcal{N}(0,1)$; otherwise it is zero. The entries are independent, and the diagonal is zero. Pairing independent entries gives $\mathbb{E}[(u^\top c)^2] = C^2 S^2 (S-1)$; the centering terms in $q_c$ contribute only $O(S^2)$ at fixed $C > 0$. Thus, for large $S$,

$$\mathbb{E}[q_c^2] = C^2 S^3 \left[1 + O(S^{-1})\right]. \tag{S63}$$

Substituting into Eq. (S62) and using $r = \sigma\sqrt{SC}$ gives

$$\frac{\mathbb{E}[\mathcal{H}_4^*]}{d} = \frac{1}{16S} \left(\frac{r}{d}\right)^4 \left[1 + O(S^{-1})\right], \qquad r = \sigma\sqrt{SC}. \tag{S64}$$

The leading large-$S$ term gives the analytical curve in Fig. 3(a) of the main text.

#### SVI.3. Elliptic Gaussian ensemble

For the fully connected elliptic Gaussian ensemble [5], distinct reciprocal pairs are independent, and the off-diagonal entries of $B$ have zero mean, unit variance, and reciprocal correlation

$$\rho = \mathrm{Corr}(B_{ij}, B_{ji}). \tag{S65}$$

To compute the moments, let $J = I_S - \mathbf{1}\mathbf{1}^\top/S$, $a = S - 1 - \rho$, and $b = (S-1)\rho - 1$. The centered row and column sums have ensemble covariance matrices

$$\mathrm{Cov}(Ju) = \mathrm{Cov}(Jc) = aJ, \qquad \mathrm{Cov}(Ju, Jc) = bJ. \tag{S66}$$

Since $q_c = (Ju)^\top (Jc)$, Gaussian fourth moments and $J^2 = J$, $\mathrm{tr}J = S-1$, give $\mathbb{E}[q_c] = (S-1)b$ and $\mathrm{Var}(q_c) = (S-1)(a^2+b^2)$. Hence

$$\mathbb{E}[q_c^2] = (S-1)(S-1-\rho)^2 + S(S-1)\big((S-1)\rho - 1\big)^2. \tag{S67}$$

Substitution into Eq. (S62) gives the finite-$S$ ensemble mean of the quartic term

$$\mathbb{E}[\mathcal{H}_4^*] = \frac{\sigma^4}{16S^2d^3}\left[(S-1)(S-1-\rho)^2 + S(S-1)\big((S-1)\rho-1\big)^2\right]. \tag{S68}$$

With $r = \sigma\sqrt{S}$, expansion at large richness gives

$$\frac{\mathbb{E}[\mathcal{H}_4^*]}{d} = \frac{1}{16}\left(\frac{r}{d}\right)^4 \left[\rho^2 + \frac{(1+\rho)(1-3\rho)}{S} + O(S^{-2})\right]. \tag{S69}$$

Within the leading weak-interaction, large-$S$ expansion, the first finite-size coefficient is positive for $-1 < \rho < 1/3$, vanishes at $\rho = 1/3$, and is negative for $\rho > 1/3$. Consequently, for sufficiently large $S$, increasing richness decreases $\mathbb{E}[\mathcal{H}_4^*]$ for $-1 < \rho < 1/3$ but increases it for $\rho > 1/3$. The theory curves in Fig. 3(b)–(d) use the complete finite-$S$ quartic ensemble mean in Eq. (S68).

#### SVI.4. Stability comparison

For the elliptic ensemble, let $\alpha(A) = \max_k \mathrm{Re}\lambda_k(A)$ be the spectral abscissa and $r = \sigma\sqrt{S}$. The elliptic law gives the local-stability margin [5, 6]

$$-\alpha(A) \approx d - r(1+\rho). \tag{S70}$$

This is the stability curve in main-text Fig. 3(b). Negative reciprocity increases this margin, while the leading large-$S$ heterogeneity depends on $\rho^2$.

### SVII. NUMERICAL AND MODEL DETAILS

This section documents the numerical procedures and model definitions used for the figures. Main-text Figs. 2 and 3 were computed in Python with NumPy and SciPy: continuous-time Lyapunov equations were solved with `scipy.linalg.solve_continuous_lyapunov`, and metapopulation equilibria with `scipy.optimize.root` (MINPACK hybrid method). Random numbers for these figures were drawn with NumPy's default generator (PCG64). The sampled-data benchmark of Fig. S1 was computed in MATLAB, as described in Sec. SIV SIV.1.

#### SVII.1. Numerical evaluation of spectral divergences

For the ecological examples in main-text Fig. 2, continuous-frequency integrals over $[0, \infty)$ were evaluated using

$$\omega = \frac{x}{1-x}, \qquad \mathrm{d}\omega = \frac{\mathrm{d}x}{(1-x)^2}, \qquad x \in [0,1). \tag{S71}$$

Midpoint quadrature used nodes $x_k = (k - 1/2)/N_\omega$, $k = 1, \ldots, N_\omega$, with equal weights $1/N_\omega$. Scalar calculations used $N_\omega = 800$ frequency points. The diamond food-web vector calculation used $N_\omega = 1200$, and the matrix divergence was evaluated from the generalized eigenvalues of the Hermitian pair $(S_Y(\omega), S_{\mathrm{OU}}(\omega))$, retaining the full complex cross-spectral matrices, including their imaginary parts. The loss integrand $\mu - \log\mu - 1$ was evaluated as $\epsilon - \log(1+\epsilon)$, with $\epsilon = \mu - 1$ and $\log(1+\epsilon)$ computed by `log1p`, to avoid cancellation when $\mu$ is close to one. The random-number generator for these ecological examples was seeded with 1.

The random-matrix calculations in main-text Fig. 3 used a 96-point Gauss–Legendre rule on $x \in (0,1)$ after the transformation

$$\omega = d\tan\Big(\frac{\pi x}{2}\Big), \qquad \mathrm{d}\omega = \frac{\pi d}{2}\sec^2\Big(\frac{\pi x}{2}\Big)\,\mathrm{d}x. \tag{S72}$$

For $\Sigma = I$ and total biomass $p = \mathbf{1}$, the exact scalar spectrum was evaluated as

$$S_Y(\omega) = \Big\|(-i\omega I - A^\top)^{-1}p\Big\|_2^2. \tag{S73}$$

For each matrix, an eigendecomposition $A^\top = W\Lambda W^{-1}$, with $\Lambda = \mathrm{diag}(\lambda_1,\dots,\lambda_n)$, was computed once and reused for all frequency nodes. With $c = W^{-1}p$, the spectrum is $S_Y(\omega) = \|W(-i\omega I - \Lambda)^{-1}c\|_2^2$, and the aggregate variance entering $\hat{\lambda}$ is

$$\gamma_Y(0) = \int_0^\infty \|e^{A^\top t}p\|_2^2\,\mathrm{d}t = \sum_{j,k}\frac{\bar{c}_j\,c_k\,(W^{\mathsf{H}}W)_{jk}}{-(\bar{\lambda}_j + \lambda_k)}, \tag{S74}$$

where $W^{\mathsf{H}}$ is the conjugate transpose. The same evaluation was used for the scalar aggregates in main-text Fig. 2(a,b). The spectral abscissa was obtained from the same eigenvalues. Figure 3 used the stability threshold $\max_k \mathrm{Re}\lambda_k(A) < -10^{-8}$; only realizations satisfying this threshold were retained. The interaction parameters of each realization ($r/d$, $\rho$, or $S$) were drawn from a generator seeded with 20260721. Each matrix was generated from its own independent stream, obtained by spawning child seeds from `numpy.random.SeedSequence(20260721)`, so the results do not depend on how realizations are distributed across parallel worker processes.

#### SVII.2. Two-species competition: main-text Fig. 2(a)

The community matrix is

$$A(\delta) = \begin{pmatrix} -1 & -0.5+\delta \\ -0.5-\delta & -1 \end{pmatrix}, \tag{S75}$$

with $\Sigma = I_2$ and total biomass $Y = X_1 + X_2$. The figure uses 41 evenly spaced values of $\delta \in [-0.45, 0.45]$, throughout which the system is Hurwitz stable; the column sums coincide at $\delta = 0$.

#### SVII.3. Spatial metapopulation: main-text Fig. 2(b)

The nonlinear metapopulation model is

$$\dot{N}_i = rN_i\left(1 - \frac{N_i}{K_i}\right) + \sum_{j\neq i} d_{ij}N_j - \sum_{j\neq i} d_{ji}N_i, \qquad i = 1,\dots,n, \tag{S76}$$

Here $N_i$ is abundance in patch $i$, $r$ the intrinsic growth rate, $K_i$ its carrying capacity, and $d_{ij}$ the dispersal rate from patch $j$ to patch $i$. Total abundance is $\mathbf{1}^\top N$. The spectral calculation uses deviations $X = N - N^*$ about the positive equilibrium $N^*$, with aggregate fluctuation $Y = \mathbf{1}^\top X$.

Patch locations are generated by a clustered point process. We sample $n_{\text{parents}} = 6$ parent points uniformly in $[0,1]^2$, then place six offspring patches around each parent using isotropic Gaussian displacements with standard deviation $s$. The total number of patches is $n = 36$, and $s$ ranges over 10 evenly spaced values in $[0.02, 0.26]$. Offspring patches falling outside $[0,1]^2$ are retained, and Euclidean distances are computed in the ambient plane.

Carrying capacities are drawn independently as

$$K_i = \max(K_{\min}, K_{\text{mean}} + K_{\text{sd}} Z_i), \qquad Z_i \sim \mathcal{N}(0,1), \tag{S77}$$

with $K_{\text{mean}} = 1.0$, $K_{\text{sd}} = 0.18$, and $K_{\min} = 0.15$.

Let $x_i \in \mathbb{R}^2$ be the location of patch $i$. Distance-dependent dispersal rates are

$$d_{ij} = m \exp\left(-\frac{\|x_i - x_j\|}{\ell}\right), \qquad i \neq j, \tag{S78}$$

where $m = 0.18$ is the dispersal amplitude and $\ell = 0.18$ the distance-decay length. The dispersal matrix $D$ has off-diagonal entries $D_{ij} = d_{ij}$ and diagonal entries $D_{ii} = -\sum_{k\neq i} d_{ki}$, so dispersal is column-conservative: $\mathbf{1}^\top D = 0$.

For each landscape realization, the positive equilibrium $N^*$ is computed with `scipy.optimize.root` (MINPACK hybrid method), using the analytical Jacobian and a relative step tolerance of $10^{-11}$. The initial guess is $N_i = K_i$; if this fails, a second initial condition $N_0 = 0.8K + 0.1U$, with $U_i \sim \text{Unif}(0,1)$, is used. A realization is accepted only if all entries of $N^*$ are positive, the residual norm is below $10^{-7}$, and the Jacobian is Hurwitz stable with threshold $\max_k \text{Re}\lambda_k(A) < -10^{-8}$.

The Jacobian at equilibrium is

$$A = D + \text{diag}\left[r\left(1 - \frac{2N_i^*}{K_i}\right)\right], \tag{S79}$$

with $r = 1$. The noise covariance is $\Sigma = I_n$. For each value of $s$, $n_{\text{rep}} = 1000$ independent landscape realizations were attempted; all 1000 satisfied the acceptance criteria at every sampled value of $s$. The plotted value is their mean, and error bars show standard errors of the mean. The insets show the first realization at $s = 0.02$, $0.10$, and $0.26$ (circled points), with patches colored by $K_i/K_{\text{mean}}$.

#### SVII.4. Diamond food web: main-text Fig. 2(c)

The diamond food web consists of a basal resource, two consumers, and a top predator, in that order. The state $X = (X_1, X_2, X_3, X_4)^\top$ contains their abundance deviations about equilibrium. The linearized dynamics have Jacobian

$$A(\delta) = \begin{pmatrix} -d & -(\alpha+\delta) & -(\alpha-\delta) & 0 \\ e(\alpha+\delta) & -d & 0 & -\beta \\ e(\alpha-\delta) & 0 & -d & -\beta \\ 0 & e\beta & e\beta & -d \end{pmatrix}, \tag{S80}$$

where $\alpha$ is the base consumption rate, $\delta$ gives equal and opposite changes in the two resource–consumer interaction strengths $\alpha \pm \delta$, $\beta$ is the predation rate, $e$ the conversion efficiency, and $d$ the self-regulation rate. Parameters are $\alpha = \beta = 0.3$, $e = 0.5$, and $d = 1$.

The coarse-graining groups species into three trophic compartments:

$$P = \begin{pmatrix} 1 & 0 & 0 & 0 \\ 0 & 1 & 1 & 0 \\ 0 & 0 & 0 & 1 \end{pmatrix}, \qquad \mathbf{Y} = (X_1, X_2 + X_3, X_4)^\top. \tag{S81}$$

The condition $PA = LP$ reduces to equality of columns 2 and 3 of $PA$, which holds if and only if $\delta = 0$. The noise covariance is $\Sigma = I_4$, and $\delta$ is swept over 41 evenly spaced values in $[0, 0.28]$.

#### SVII.5. Random community matrices: main-text Fig. 3

Main-text Fig. 3 shows individual stable realizations together with kernel estimates of their ensemble mean and of their 10–90% range, described at the end of this section. The counts below refer to attempted matrices; only realizations satisfying the stability threshold in Sec. SVII SVII.1 and yielding finite positive numerical losses are retained. For the parameters used, all attempted realizations were retained. In every panel, $d = 1$, $\Sigma = I_S$, and total biomass is $Y = \mathbf{1}^\top X$. The interaction parameter for each realization is drawn independently from the indicated range. Theory curves show the quartic ensemble mean $\mathbb{E}[\mathcal{H}_4^*]$ derived in Sec. SVI.

*Interaction-strength sweep: Fig. 3(a)*

Each off-diagonal entry $A_{ij}$, $i \neq j$, is independently drawn from $\mathcal{N}(0, \sigma^2)$ with probability $C$ and is zero with probability $1 - C$; diagonal entries are $-d$. Parameters are $S = 100$, $C = 0.3$, and $d = 1$. For each of 6000 independent realizations, the dimensionless interaction strength $x = r/d$ is sampled uniformly on the linear interval $[0.05, 0.80]$, and

$$\sigma = \frac{xd}{\sqrt{SC}}.$$

The dashed curve uses Eq. (S64). It is a weak-interaction approximation; higher-order contributions need not be small at the upper end of this sweep.

*Reciprocal-correlation sweep: Fig. 3(b)*

Community matrices have prescribed pairwise correlation $\rho = \mathrm{Corr}(A_{ij}, A_{ji})$. For each pair $i > j$,

$$A_{ij} = \sigma Z_1, \qquad A_{ji} = \sigma \left( \rho Z_1 + \sqrt{1 - \rho^2}\, Z_2 \right), \tag{S82}$$

where $Z_1, Z_2 \sim \mathcal{N}(0, 1)$ are independent. The network is fully connected, with $S = 100$, $d = 1$, and fixed $r/d = 0.15$, so

$$\sigma = \frac{r}{\sqrt{S}} = 0.015.$$

For each of 7500 independent attempted realizations, $\rho$ is sampled uniformly on $[-0.9, 0.9]$. The heterogeneity curve uses the complete finite-$S$ quartic ensemble mean in Eq. (S68). The local-stability curve uses Eq. (S70).

*Richness sweeps: Figs. 3(c) and 3(d)*

The richness sweeps use the fully connected elliptic ensemble with $d = 1$ and fixed $r/d = 0.15$. Species richness is sampled uniformly as an integer on $\{50, 51, \ldots, 500\}$, and

$$\sigma = \frac{r}{\sqrt{S}}$$

is adjusted for each realization to keep the collective interaction scale fixed. Panel (c) uses $\rho = 0$, and panel (d) uses $\rho = 0.8$. For each panel, 7500 independent realizations are attempted. The dashed curves use the complete finite-$S$ quartic ensemble mean in Eq. (S68); the dotted line in panel (d) is its large-$S$ limit, $\rho^2(r/d)^4/16$.

*Ensemble-mean and spread estimates*

The solid curves in main-text Fig. 3 estimate the conditional ensemble mean $\mathbb{E}[\mathcal{H}^* \mid x]$, where $x$ is $r/d$ in panel (a), $\rho$ in panel (b), and $S$ in panels (c) and (d). Because $\mathcal{H}^*$ spans several decades along $x$, smoothing it directly would bias the estimate. We therefore smooth the ratio $z_i = \mathcal{H}_i^*/T(x_i)$, where $T(x)$ is the analytical curve shown in the same panel, and report $T(x)\,\hat{\mathbb{E}}[z \mid x]$. Since $T(x)$ is deterministic, $T(x)\,\mathbb{E}[z \mid x] = \mathbb{E}[\mathcal{H}^* \mid x]$ exactly; the normalization changes only the smoothing bias. The estimate $\hat{\mathbb{E}}[z \mid x]$ is a local-linear regression with Gaussian kernel weights

$$w_i(u) = \exp\left[-\frac{(u_i - u)^2}{2h^2}\right] \tag{S83}$$

in the plotted coordinate $u$: $u = \log_{10}(r/d)$ in panel (a), $u = \rho$ in panel (b), and $u = \log_{10} S$ in panels (c) and (d), with bandwidths $h = 0.05$, 0.05, and 0.08, respectively.

The shaded bands show the 10% and 90% quantiles of $\mathcal{H}^*$ conditional on $x$, obtained as $T(x)$ times the kernel-weighted empirical quantiles of $z$; this is exact for quantiles because $T(x) > 0$. Panel (a) uses $h = 0.1$ for the band. The band is drawn only where the effective sample size $\left(\sum_i w_i\right)^2 / \sum_i w_i^2$ is at least 300; below this, tail quantiles of the strongly skewed distribution of $\mathcal{H}^*$ are too noisy on a logarithmic axis. This criterion removes only $r/d \lesssim 0.06$ in panel (a). The bands describe the spread of individual realizations, not the uncertainty of the mean; with 6000–7500 realizations per panel, 95% bootstrap confidence intervals of the mean are not resolvable at the scale of the plots over most of each panel.

*Simulated scalings*

To compare simulated and predicted trends quantitatively, we fitted mean models with variance proportional to the squared mean (Gamma quasi-likelihood, fitted by iteratively reweighted least squares) and obtained 95% intervals from 2000 bootstrap resamples of realizations. In panel (a), restricted to $r/d \leq 0.3$, the fit $\mathbb{E}[\mathcal{H}^*] \propto (r/d)^\beta$ gives $\beta = 4.05$ $[3.90, 4.19]$, consistent with the quartic prediction; above $r/d \approx 0.4$, the simulated mean exceeds the quartic term, as expected from higher-order contributions. In panel (c), $\mathbb{E}[\mathcal{H}^*] \propto S^\beta$ gives $\beta = -1.05$ $[-1.11, -1.00]$, compared with $-0.98$ for the finite-$S$ prediction over $50 \leq S \leq 500$. In panel (d), the fit

$$\frac{16\,\mathbb{E}[\mathcal{H}^*]}{d\,(r/d)^4} = A + \frac{B}{S} \tag{S84}$$

gives $A = 0.689$ $[0.683, 0.695]$ and $B = -4.3$ $[-5.6, -3.0]$, with $B < 0$ in all bootstrap resamples, compared with the predictions $A = \rho^2 = 0.64$ and $B = (1+\rho)(1-3\rho) = -2.52$. The simulations thus confirm the predicted increase of the mean with richness at $\rho = 0.8$, while the larger simulated level and finite-size coefficient indicate higher-order corrections at $r/d = 0.15$. In panel (b), the ratio of the simulated mean to Eq. (S68) is $0.99 \pm 0.02$ for $-0.9 \leq \rho \leq -0.5$ and $1.05 \pm 0.02$ for $0.5 \leq \rho \leq 0.9$ (mean $\pm$ 95% interval), so the leading-order symmetry under $\rho \to -\rho$ holds to within about 5% at this interaction strength.

---

[1] F. Itakura and S. Saito, Analysis synthesis telephony based on the maximum likelihood method, in *Proceedings of the 6th International Congress on Acoustics* (1968) pp. C17–C20.

[2] A. Ferrante, C. Masiero, and M. Pavon, Time and spectral domain relative entropy: A new approach to multivariate spectral estimation, IEEE Transactions on Automatic Control **57**, 2561 (2012).

[3] Y. Iwasa, V. Andreasen, and S. A. Levin, Aggregation in model ecosystems. I. Perfect aggregation, Ecol. Modelling **37**, 287 (1987).

[4] R. M. May, Will a large complex system be stable?, Nature **238**, 413 (1972).

[5] H.-J. Sommers, A. Crisanti, H. Sompolinsky, and Y. Stein, Spectrum of large random asymmetric matrices, Phys. Rev. Lett. **60**, 1895 (1988).

[6] S. Allesina and S. Tang, Stability criteria for complex ecosystems, Nature **483**, 205 (2012).
